# H3K4 di- and trimethylation modulate the stability of RNA polymerase II pausing

**DOI:** 10.1101/2022.11.28.518073

**Authors:** Shibin Hu, Aixia Song, Linna Peng, Nan Tang, Zhibin Qiao, Zhenning Wang, Fei Xavier Chen

## Abstract

Modifications of histones are intricately linked with the regulation of gene expression, with demonstrated roles in various physiological processes and disease pathogenesis. Methylation of histone H3 lysine 4 (H3K4), implemented by the COMPASS family, is enriched at promoters and associated cis-regulatory elements, with H3K4 trimethylation (H3K4me3) considered a hallmark of active gene promoters. However, the relative roles of deposition and removal of H3K4 methylation, as well as the extent to which these events contribute to transcriptional regulation have so far remained unclear. Here, through rapid depletion of the transcription regulator SPT5 or either of two shared subunits of COMPASS family members, we reveal a dynamic turnover of H3K4me3 mediated by the KDM5 family of histone demethylases. Loss of H3K4me3 following COMPASS disruption does not impair the recruitment of TFIID and initiating RNA polymerase II (Pol II). Instead, H3K4me3 loss leads to reductions in the paused form of Pol II on chromatin while inducing the relative enrichment of the Integrator-PP2A (INTAC) termination complex, leading to reduced levels of elongating polymerases, thus revealing how H3K4me3 dynamics can regulate Pol II pausing to sustain or attenuate transcription.

## INTRODUCTION

For either protein-coding genes or non-coding elements, transcription is driven by DNA-binding transcription factors that recognize specific DNA motifs to guide the landing of the transcription machinery at promoters and its subsequent progression of it through gene bodies (*1–3*). Eukaryotic DNA, along with histone octamers, is wrapped into strings of nucleosomes to assemble a structure called chromatin, which serves as a versatile platform enabling the complex communications between transcription regulators and genetic information. Histones are subject to a variety of posttranslational modifications, plenty of which are heritable and thus are considered epigenetic marks. These marks not only influence chromatin compaction and nucleosome dynamics, but also directly modulate the binding of many chromatin and transcription regulators harboring “reader” domains for their corresponding marks. As such, dynamic regulation of these marks by a group of writers and erasers is a fundamental mechanism for achieving precise control of gene expression, while dysregulation of these processes is frequently observed in developmental disorders and cancers (*4–14*).

Among the most intensively studied epigenetic marks, histone H3 lysine 4 (H3K4) methylation is evolutionally conserved and solely depends on Set1/COMPASS for its deposition. H3K4 trimethylation (H3K4me3) is consistently found around transcription start sites (TSSs) and serves as a hallmark of active gene promoters. In contrast, H3K4me1 is prevalent at intragenic and intergenic cis-regulatory elements with a relative enrichment at enhancer regions and, thus, has been widely used as a marker for active or poised enhancers. The COMPASS family in mammals comprises six members: SET1A, SET1B, MLL1, MLL2, MLL3, and MLL4, each of which participates in similar complexes containing four shared subunits collectively known as WRAD (WDR5, RBBP5, ASH2L, and DPY30) (*15–20*). Despite having some level of redundancy, COMPASS family members differ in their product specificity: SET1A and SET1B are primarily responsible for H3K4me3 at promoters; MLL1 and MLL2 mainly catalyze H3K4me2 with selective H3K4me3 activity at specific loci, such as “bivalently marked” chromatin in embryonic stem cells (ESCs); while MLL3 and MLL4 predominantly implement H3K4me1 at enhancers (*21–29*). However, it is important to note that, individual perturbation of one COMPASS member can be partially compensated by other family members. For example, individual ablation of SET1A or SET1B in mouse ESCs does not perturb global H3K4me3 levels, while simultaneous disruption of SET1A and SET1B causes a limited decline in bulk levels of H3K4me3 (*30*). For this reason, it has been challenging to achieve a near-complete loss of H3K4 methylation in cells, especially in mammals, which has hindered the elucidation the full range of functions of these histone modifications.

The removal of H3K4 methylation is catalyzed by two families of histone demethylases, the KDM1 family (KDM1A and KDM1B) that is only capable of demethylating the mono and dimethylated H3K4 and the KDM5 family (KDM5A, KDM5B, KDM5C, and KDM5D) that preferentially targets H3K4me3 and H3K4me2 (*31–34*). However, the degree to which these marks are relatively stable or dynamically regulated, and how the relevant methyltransferases and demethylases cooperate to control the kinetics for each of the three methylation states for the regulation of transcription or other DNA-templated processes are important yet outstanding questions.

Despite its high correlation with active transcription, the instructive role of H3K4me3 in transcriptional activation is less well understood and even debated (*35, 36*). H3K4me3 potentiates or modulates gene expression by facilitating the recruitment of specific methyl-binding proteins, including TAF3, CHD1, BPTF, SGF29, ING2, ING5, BAP18, and SPIN1/2/3 (*37–42*). Among them, the interaction between H3K4me3 and TAF3, a crucial subunit of the general transcription factor TFIID, is arguably the most well-known, as TFIID is crucial for establishing transcription initiation at promoters, regions which have the highest levels of H3K4me3. Therefore, several landmark studies have led to a model where H3K4me3 is important for TFIID recruitment and/or stabilization and, thus, facilitates the assembly of the transcription preinitiation complex (PIC) (*38, 39, 43-45*). Nonetheless, an instructive role of H3K4me3 has also been challenged as some perturbations of this modification had a limited impact on gene expression, especially in S. cerevisiae (*35, 36*). It is unclear whether this disparity between species is owing to differences in reader proteins (e. g., yeast TAF3 does not contain a PHD that recognizes H3K4me3) or distinctions in transcriptional regulation mechanisms, such as promoter-proximal pausing, which is a pervasive feature of metazoan gene regulation (*2, 3, 46*).

Here, by using acute endogenous protein degradation strategies, we reveal that, upon the loss of SPT5 or disruption of COMPASS, H3K4me3 is rapidly removed in a KDM5 demethylase-dependent manner. Surprisingly, large scale reductions in H3K4me2 and H3K4me3 exerted no appreciable effects on the genome-wide recruitment of TFIID and initiating polymerases. However, levels of paused Pol II appreciably decreased, while relative levels of the pausing regulator INTAC increased following the reduction of H3K4me2 and H3K4me3. Therefore, our results uncover a dynamic regulation of H3K4 methylation and its role in regulating Pol II pausing.

## RESULTS

### SPT5 regulates global H3K4 methylation

We previously reported that the acute depletion of SPT5, a pleiotropic transcription regulator that governs multiple transcription steps from pausing to termination, influences the openness of a subset (∼5%) of active promoters and enhancers (*47*). Consequently, bulk levels of histone H3 lysine 27 acetylation (H3K27ac), a chromatin mark found at promoters and enhancers, remain unchanged upon the degradation tag FKBP12^F36V^ (dTAG)-mediated acute degradation of SPT5 in human DLD-1 cells (Fig. 1A-B). To determine whether SPT5 regulates other active transcription marks, we measured the bulk levels of H3 lysine 4 (H3K4) mono-(H3K4me1), di-(H3K4me2), and trimethylation (H3K4me3) upon SPT5 depletion. Surprisingly, 6-hour dTAG treatment leads to a marked decrease in H3K4me3 levels but not H3K4me2 or H3K4me1, in contrast with the previous notion that H3K4me3 is a relatively long-lived histone modification and could potentially act as a heritable “memory” mark of active transcription (*48*) (Fig. 1B). We conducted ChIP-seq with reference exogenous genome as the spike-in (ChIP-Rx) (*49*) for H3K4me3 and H3K27ac. In line with the western blotting results, the levels of H3K4me3, but not H3K27ac, decline dramatically at promoter regions by 6-hour SPT5 depletion (Fig. 1C-D, Fig. S1A-B).

**Figure 1.**
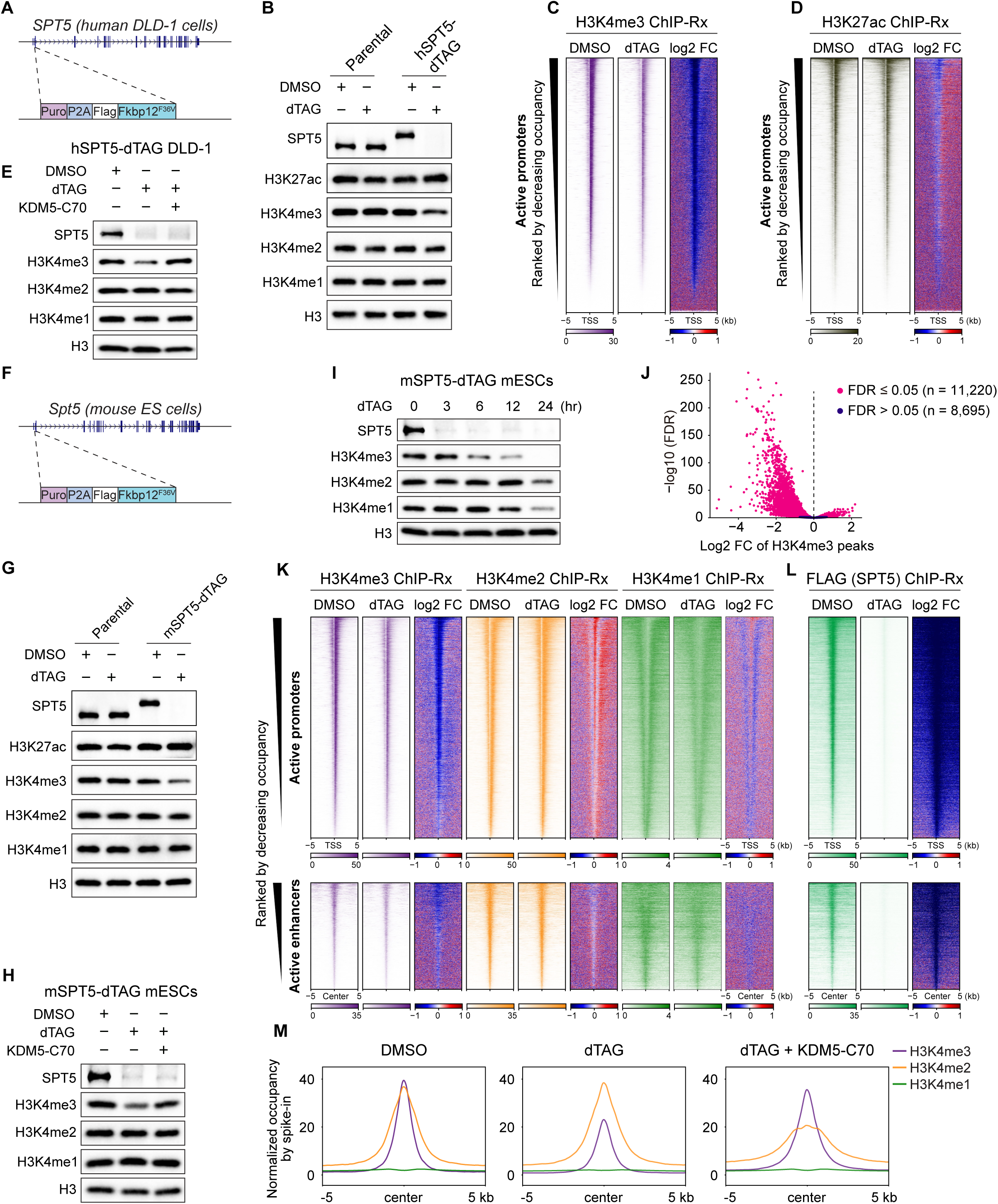
SPT5 depletion induces a global loss of H3K4me3. (A) Schematic of the generation of knock-in hSPT5-dTAG DLD-1 cells. (B) Western blotting of whole-cell extracts of parental and hSPT5-dTAG DLD-1 cells treated with DMSO or dTAG for 6 hours. Histone H3 is a loading control. (C and D) Heatmaps of H3K4me3 (C) and H3K27ac (D) occupancies (reads per million [RPM] per base pair [bp] and log2 fold change) of hSPT5-dTAG DLD-1 cells centered at TSS of promoters ranked by decreasing occupancy in the DMSO condition. (E) Western blotting of dTAG treatment with or without pan-KDM5 inhibition by KDM5-C70 in hSPT5-dTAG DLD-1 cells. (F) Schematic of the generation of mSPT5-dTAG mouse ES cells (mESCs) by knock-in. (G) Western blotting of whole-cell extracts of parental and mSPT5-dTAG mESCs treated with DMSO or dTAG for 6 hours. H3 is a loading control. (H) Western blotting of dTAG treatment with or without pan-KDM5 inhibition by KDM5-C70 in mSPT5-dTAG mESCs. (I) Western blotting of mSPT5-dTAG mESCs with time-course dTAG treatment. (J) Volcano plot showing the log2 fold change of H3K4me3 peaks of 6-hour dTAG versus DMSO treatment. (K) Heatmaps of H3K4me3, H3K4me2, and H3K4me1 occupancies (RPM per bp and log2 fold change) ranked by decreasing H3K4me3 occupancy in the DMSO condition. The peaks are centered at the TSS of genes with H3K4me3 promoter peaks for active promoters (top), and at H3K4me3 peak centers for active enhancers (bottom). (L) Heatmaps of FLAG (SPT5) occupancy (RPM per bp and log2 fold change) ranked by decreasing H3K4me3 occupancy in the DMSO condition. The peaks are centered at the same positions as (K). (M) Metaplots showing the average levels of H3K4me3, H3K4me2, and H3K4me1 occupancy centered at H3K4me3 peak centers in mSPT5-dTAG mESCs treated with DMSO, dTAG, or dTAG + KDM5-C70. See also Figure S1.

H3K4me3 demethylation is catalyzed by members of the KDM5 family (*50, 51*). Treatment with KDM5-C70, a pan-KDM5 inhibitor, restores H3K4me3 levels in dTAG-treated cells, indicating that the KDM5 family is responsible for the rapid removal of H3K4me3 in SPT5-depleted cells (Fig. 1E). To confirm that the rapid decline in H3K4me3 is not specific to cancer cell lines, we integrated dTAG into the N-terminus of the endogenous *Spt5* locus in mouse embryonic stem cells (mESCs), which is denoted as mSPT5-dTAG cells (Fig. 1F). Consistently, 6-hour dTAG treatment induces a near-complete depletion of SPT5 and a prominent decline in H3K4me3 in mESCs (Fig. 1G), with the latter being rescued by KDM5-C70 treatment (Fig. 1H).

We next assessed the dynamics of H3K4 methylation by conducting a time course of SPT5 degradation in mSPT5-dTAG cells. The observed decrease in H3K4me3 is faster than that of H3K4me2 or H3K4me1 following dTAG treatment (Fig. 1I), consistent with previous studies showing that KDM5 is more active in demethylating H3K4me3 (*52–57*). We conducted ChIP-Rx in mSPT5-dTAG cells treated with DMSO or dTAG for 6 hours to map the genome-wide alterations of all three H3K4 methylation states. Among all 19,945 H3K4me3 peaks, a majority of significantly changed peaks exhibit decreased H3K4me3 levels (Fig. 1J, Fig. S1C). We thus divided all H3K4me3 peaks into two groups: significantly decreased and non-decreased peaks. In contrast to the pronounced reduction in H3K4me3 for around half of the peaks, levels of H3K4me1 and H3K4me2 are only slightly affected at these regions (Fig. S1D), consistent with the western blotting results (Fig. 1G). Analysis of peak positions reveals that 89.1% of significantly decreased H3K4me3 peaks are at promoter regions, while the non-decreased peaks have 66.4% overlap with promoters and 33.6% overlap with enhancers (Fig. S1E-F). Indeed, the decline in H3K4me3 is more apparent at promoters than at enhancers (Fig. 1K). Moreover, the Pol II levels at promoters measured by precision run-on sequencing (PRO-seq) are markedly higher for the H3K4me3 decreased group than the non-decreased group (Fig. S1G). The loss of SPT5 is equally evident at all H3K4me3-covered loci regardless of their classification as promoters or enhancers, indicating that greater H3K4me3 changes at promoters resulting from dTAG treatment is not a result of differences in SPT5 degradation efficiency at specific loci (Fig. 1L, Fig. S1D). Collectively, these data suggest that SPT5 preferentially regulates H3K4me3 levels at active promoters having higher levels of paused Pol II.

To examine the genome-wide impact of the KDM5 family on SPT5 loss-induced H3K4me3 reductions, we conducted ChIP-Rx of H3K4 methylation with or without KDM5 inhibition. The resulting analysis demonstrates that decreased H3K4me3 levels resulting from SPT5 depletion are largely restored by KDM5-C70 treatment (Fig. 1M, purple lines). Interestingly, although 6 hours of dTAG treatment alone does not evidently affect H3K4me2 levels, the addition of KDM5-C70 results in a noticeable reduction in H3K4me2 levels, presumably through blocking the KDM5-mediated conversion of H3K4me3 to H3K4me2 (Fig. 1M, orange lines). Taken together, we conclude that, at least in the human DLD-1 cells and mESCs we tested, H3K4me3 is dynamically regulated and its removal depends on the KDM5 family of demethylases.

### COMPASS continuously maintains H3K4 methylation

Deposition of methylation on H3K4 is catalyzed by members of the COMPASS (complex of proteins associated with Set1) family, including SET1A, SET1B, and MLL1-4. Although SET1A and SET1B in mammalian cells are mainly responsible for bulk H3K4me3, MLL1 and MLL2 catalyze H3K4me3 in a locus-specific manner and can partially compensate for SET1A/SET1B loss in H3K4me3 deposition (*20*). Among the four shared COMPASS subunits known as WRAD (WDR5, RBBP5, ASH2L, and DPY30), RBBP5 is essential for complex assembly, nucleosome recognition, and thus catalytic activation, whereas DPY30 flexibly associates with and stimulates H3K4 methylation (*21, 58–65*). Therefore, to investigate the dynamics of genome-wide H3K4 methylation, we established mRBBP5-dTAG and mDPY30 mESCs to individually degrade RBBP5 or DPY30 (Fig. 2A and Fig. S2A). 6-hour dTAG treatment in mRBBP5-dTAG cells induces a substantial loss in H3K4me2 and H3K4me3, while H3K4me1 exhibits no noticeable change at the bulk level (Fig. 2B). Incubating dTAG-treated cells with KDM5-C70 restores H3K4me3 levels, indicating that the KDM5 family is mainly responsible for RBBP5 degradation-induced H3K4me3 demethylation, similarly to what was observed in SPT5-depleted cells (Fig. 2C). Time-resolved western blotting results reveal that H3K4me3 levels decline more rapidly than those of H3K4me1 and H3K4me2, with a near-complete loss of H3K4me3 achieved after 12-hour dTAG treatment (Fig. 2D).

**Figure 2.**
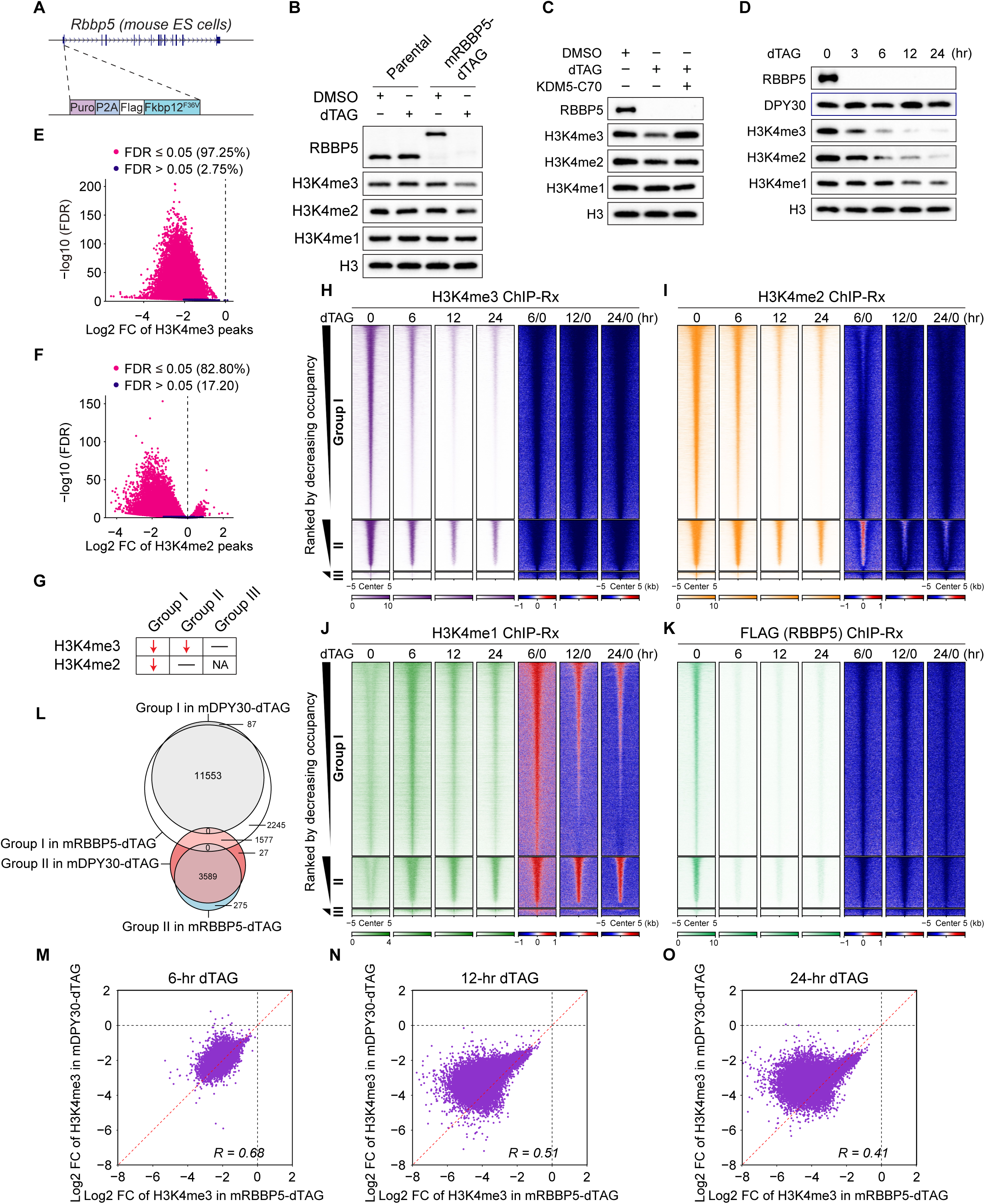
COMPASS is continuously required to maintain H3K4 methylation. (A) Schematic of the generation of knock-in mRBBP5-dTAG mESCs. (B) Western blotting of whole-cell extracts of parental and mRBBP5-dTAG mESCs treated with DMSO or dTAG for 6 hours. H3 is a loading control. (C) Western blotting of dTAG treatment with or without pan-KDM5 inhibition by KDM5-C70 in mRBBP5-dTAG mESCs. (D) Western blotting of mRBBP5-dTAG mESCs with indicated time-course of dTAG treatment. (E and F) Volcano plots showing the log2 fold change of H3K4me3 peaks (E) and H3K4me2 peaks (F) of 6-hour dTAG versus DMSO treatment. (G) Schematic showing the change in patterns of three groups of peaks for 6-hour dTAG versus DMSO treatment. (H-K) Heatmaps showing three groups of H3K4me3 (H), H3K4me2 (I), H3K4me1 (J), and FLAG (RBBP5) (K) occupancy (RPM per bp and log2 fold change) ranked by decreasing H3K4me3 occupancy in the DMSO condition. The peaks are centered at H3K4me3 peak centers. (L) Venn diagrams showing overlaps of Group I and Group II H3K4me3 peaks between mDPY30-dTAG mESCs and mRBBP5-dTAG mESCs. (M-O) Scatterplots depicting the correlations of log2 fold change (FC) of H3K4me3 peaks of 6- (M), 12- (N), and 24-hour dTAG (O) versus DMSO treatment between mRBBP5-dTAG cells and mDPY30-dTAG cells. See also Figure S2.

To monitor the dynamic alterations in H3K4 methylation genome-wide, we performed ChIP-Rx in dTAG-treated mRBBP5-dTAG mESCs at different time points. After 6 hours of dTAG treatment, all significantly changed peaks of H3K4me3, and most of the significantly changed peaks of H3K4me2, had decreased occupancy (Fig. 2E-F). We therefore classified the peaks into three groups using this time point: Group I are loci showing significant decline in both H3K4me2 and H3K4me3, Group II are loci with only H3K4me3 reduction, and Group III are loci exhibiting no significant change in H3K4me3 (Fig. 2G). For Group I peaks, both H3K4me2 and H3K4me3 levels were dramatically reduced at 6 hours and reached minimal levels by 12-24 hours (Fig. 2H-J, upper part). For Group II peaks that have higher average H3K4me3 levels in the 0-hour condition, H3K4me3 declines at a slower rate than that of Group I, while H3K4me1 and H3K4me2 reach a summit at 6 hours but gradually decrease thereafter (Fig. 2H-J, middle part). Group III contains peaks with extremely low levels of all three methylation states and was therefore excluded from subsequent analyses (Fig. 2H-J, lower part). Importantly, and in line with the western blotting results, dTAG treatment resulted in a rapid and robust depletion of RBBP5 protein for all groups of peaks, indicating that differences in H3K4 methylation changes are unlikely to result from varying RBBP5 degradation efficiencies at specific loci (Fig. 2K).

In accordance with observations in mRBBP5-dTAG mESCs, acute DPY30 degradation induced a rapid decline in H3K4me3 (Fig. S2A-B), and the addition of the KDM5 inhibitor KDM5-C70 restored H3K4me3 levels (Fig. S2C). Moreover, although the overall rate of change in H3K4 methylation states is somewhat slower over the course of dTAG treatment in mDPY30-dTAG compared with mRBBP5-dTAG mESCs, the relative kinetics of mono vs. di vs. trimethylation changes are comparable, with H3K4me3 showing the fastest decline and H3K4me1 exhibiting the slowest change (Fig. S2D). We thus used the same criteria to group H3K4me3 peaks affected by DPY30 depletion that we used for characterizing changes resulting from RBBP5 depletion (Fig. 2G). We found that the initial occupancy and the time-resolved changes in H3K4 methylation states were similar in mDPY30-dTAG and mRBBP5-dTAG cells (Fig. S2E-H).

Despite the above noted similarities, we noticed that a larger number of peaks show simultaneous loss of H3K4me2 and H3K4me3 by 6 hours of degradation of RBBP5 (n = 13,798) than 6 hours of DPY30 degradation (n = 11,640), in accordance with the fact that RBBP5 is more crucial for COMPASS functions (*58, 59, 61, 62, 66*) (Fig. 2L). To further validate this conclusion, we compared the alterations of H3K4 methylation states at all time points tested in RBBP5-dTAG and DYP30-dTAG cells. Although the changes in all three forms of H3K4 methylation are positively correlated, the reduction of H3K4me2 and H3K4me3 induced by RBBP5 depletion is more pronounced than by DPY30 depletion (Fig. 2M-O, Fig. S2I-N). In particular, H3K4me3 loss on a large number of peaks was several times greater following 24-hour dTAG treatment in mRBBP5-dTAG than in mDPY30-dTAG cells. We therefore mainly used mRBBP5-dTAG mESCs to further explore biological functions of H3K4 methylation.

### H3K4 methylation is required for maintaining Pol II levels at promoters independently of TFIID recruitment

Having established the dynamic changes of H3K4 methylation states following COMPASS disruption, we next asked how these changes impact transcription by performing time-resolved Pol II ChIP-Rx in dTAG-treated mRBBP5-dTAG mESCs. The genome-wide changes of Pol II occupancy at H3K4me3-enriched regions were analyzed separately for promoters and enhancers (Fig. S3A). Both promoters and enhancers in our previously defined Group I and II loci (Fig. 2G), exhibited reduced Pol II occupancy by 12- and 24- hour dTAG treatment, while 6 hours of dTAG treatment had no notable changes (Fig. 3A, Fig. S3B-C). Because both the alterations of H3K4 methylation and Pol II levels are equivalently evident for Groups I and II, especially at promoters, we combined the two groups and focused on promoter regions for our subsequent analysis, which confirmed the prevalent decrease in Pol II occupancy at promoters after 12 and 24 hours of dTAG treatment (Fig. 3B, Fig. S3D).

**Figure 3.**
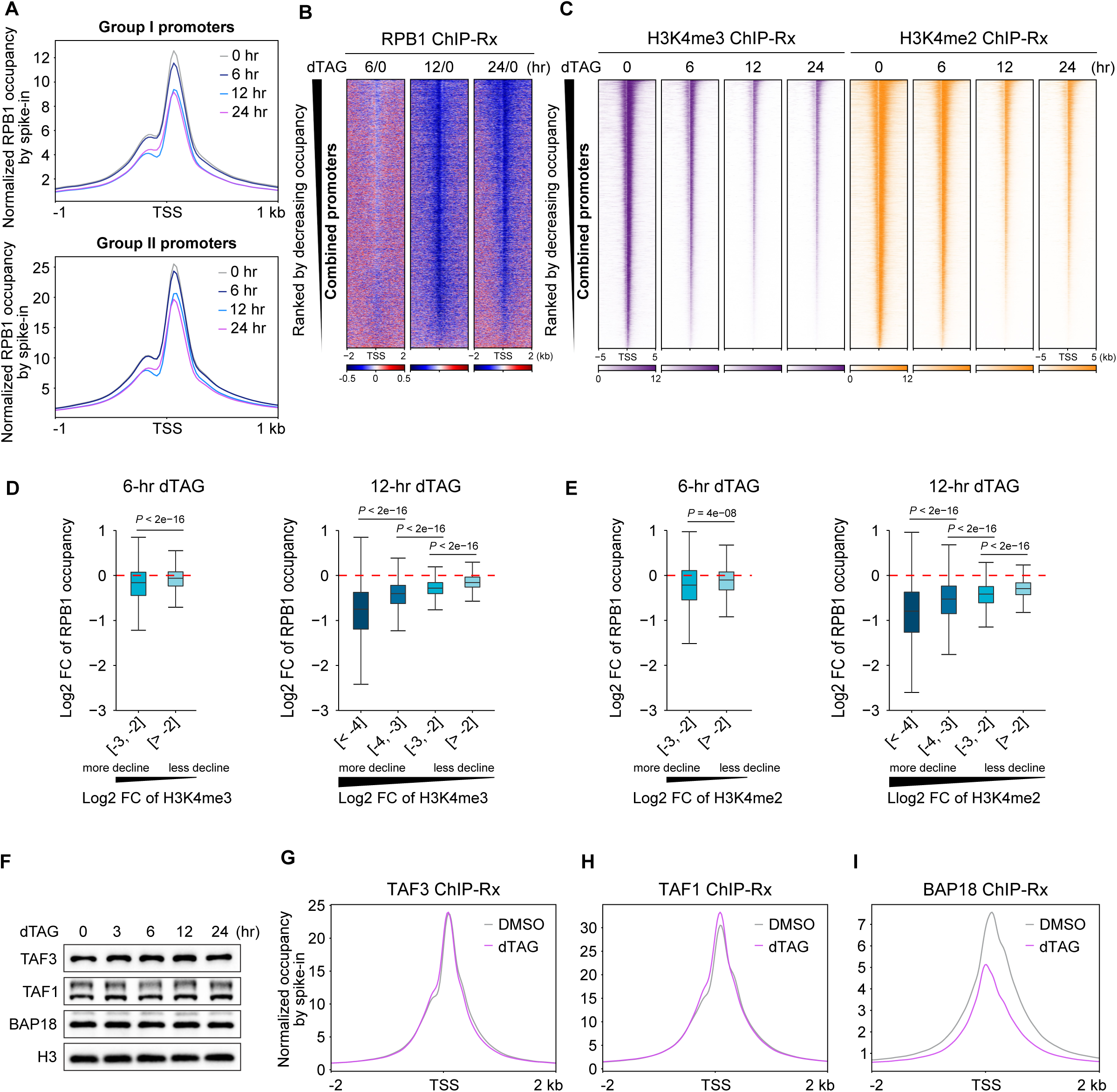
RBBP5 depletion and subsequent loss of H3K4me2 and H3K4me3 reduce Pol II levels at promoters independently of TFIID recruitment. (A) Metaplots showing the average level of Pol II occupancy at Group I promoters (top) and Group II promoters (bottom) centered at the TSS of genes with H3K4me3 promoter peaks. Pol II occupancy is represented by its largest subunit RPB1. (B) Fold-change heatmaps of Pol II occupancy (RPM per bp) (dTAG versus DMSO) in mRBBP5-dTAG cells ranked by decreasing Pol II occupancy in the DMSO condition. (C) Heatmaps showing the occupancies (RPM per bp) of H3K4me3 and H3K4me2 in mRBBP5-dTAG mESCs indicated time-course of dTAG treatment. The peaks are ranked by decreasing H3K4me3 occupancy in the DMSO condition. (D) Boxplots showing the correlation of log2 fold change of Pol II (dTAG versus DMSO) and H3K4me3 (dTAG versus DMSO, grouped based on the extent of declining H3K4me3 occupancy) in mRBBP5-dTAG mESCs. (E) Boxplots showing the correlation of log2 fold change of Pol II (dTAG versus DMSO) and H3K4me2 (dTAG versus DMSO, grouped based on the extent of declining H3K4me3 occupancy) in mRBBP5-dTAG mESCs. (F) Western blotting of mRBBP5-dTAG mESCs with indicated time-course dTAG treatment. (G-I) Metaplots showing the average levels of TAF3, TAF1, and BAP18 occupancy centered at TSS of genes with H3K4me3 promoter peaks. See also Figure S3.

Given that changes in H3K4me2 and H3K4me3 levels are first apparent at 6 hours of RBBP5 depletion, but their near depletion by 12-24-hour dTAG-treated cells (Fig. 3C, Fig. S3E-F) was concomitant with decreased Pol II occupancy, we speculated that a critical level of H3K4 methylation was required to maintain Pol II at promoters. To validate this hypothesis, we compared changes in H3K4me2 and H3K4me3 levels with Pol II occupancy at promoters after 6 and 12 hours of dTAG treatment. Notably, although decreased H3K4 methylation is positively correlated with decreased Pol II occupancy for both time points, the greatest decline in Pol II occupancy occurs after 12 hours of dTAG treatment when H3K4 methylation levels have dropped more than 16 fold (log2FC < −4) (Fig. 3D-E). The results indicate that a near-complete depletion of H3K4me2 and H3K4me3 is likely required to disrupt the recruitment and/or maintenance of Pol II at promoters.

Being a hallmark of active gene promoters, H3K4me3 has been proposed to play an instructive role in transcription by facilitating the recruitment of H3K4me3 readers or reader-containing complexes, such as the preinitiation complex (PIC) assembly factor TFIID. Therefore, we sought to determine whether the reduced Pol II occupancy at promoters in COMPASS-disrupted cells can be ascribed to impaired recruitment of TFIID. TAF3, a subunit of TFIID, which can directly interact with H3K4me3 (*38, 39, 43*), and TAF1, a scaffold subunit nucleating TFIID assembly and recognizing DNA (*67–70*), were selected for ChIP-Rx to quantify TFIID occupancy genome wide. BAP18, as another demonstrated reader protein of H3K4me3 (*37*) was included as a control. Bulk levels of examined factors are not affected by dTAG-induced RBBP5 degradation (Fig. 3F). Surprisingly, even with 24-hour dTAG-induced RBBP5 degradation and a near-complete loss of both H3K4me2 and H3K4me3, the chromatin occupancies of TAF3 or TAF1 exhibit no notable alterations compared with DMSO-treated cells (Fig. 3G-H, Fig. S3G-H). Contrastingly, BAP18 levels at promoters decline dramatically in RBBP5-depleted mESCs (Fig. 3I, Fig. S3I). We conclude that TFIID recruitment is not perturbed upon disruption of COMPASS and the subsequent loss of H3K4me2 and H3K4me3 in mESCs.

### H3K4 methylation contributes to the maintenance of paused but not initiating Pol II

Pol II ChIP-seq signal reflects a combination of non-engaged polymerases within the PIC and transcriptionally engaged but transiently paused Pol II. The transition of Pol II from the PIC to the pausing complex requires the translocase activity of XPB/TFIIH (Fig. 4A). The unexpected results that the acute RBBP5 loss exerts no notable impact on TFIID recruitment prompted us to determine which type of Pol II is primarily affected in RBBP5-depleted cells. Since the residency time of paused Pol II for most active genes is shorter than 10 minutes (min), we treated the cells with triptolide (TPL), an inhibitor of the translocase activity of XPB/TFIIH, for 10 and 20 min to completely deplete new Pol II forming the pausing complex while preserving Pol II within the PIC. As expected, TPL treatment causes a 5’ shift of Pol II peak centers (Fig. 4B). We next measured Pol II levels under these conditions in dTAG-treated RBBP5-dTAG cells. In stark contrast to the evident reduction of Pol II levels at promoters after RBBP5 degradation without TPL (Fig. 4C), dTAG treatment does not reduce promoter-bound Pol II under 10- or 20-min TPL-treated conditions (Fig. 4D-E). These data suggest that H3K4me2 and H3K4me3 loss preferentially impairs the enrichment of paused Pol II rather than PIC at promoters. Indeed, RBBP5 depletion even leads to a slight increase in levels of Pol II within PIC under TPL treatment conditions (Fig. 4D-E), as Pol II in the paused state inhibits new transcription initiation and, thus, greater loss of paused polymerase in RBBP5-depleted cells could potentially promote transcription initiation (*71, 72*).

**Figure 4.**
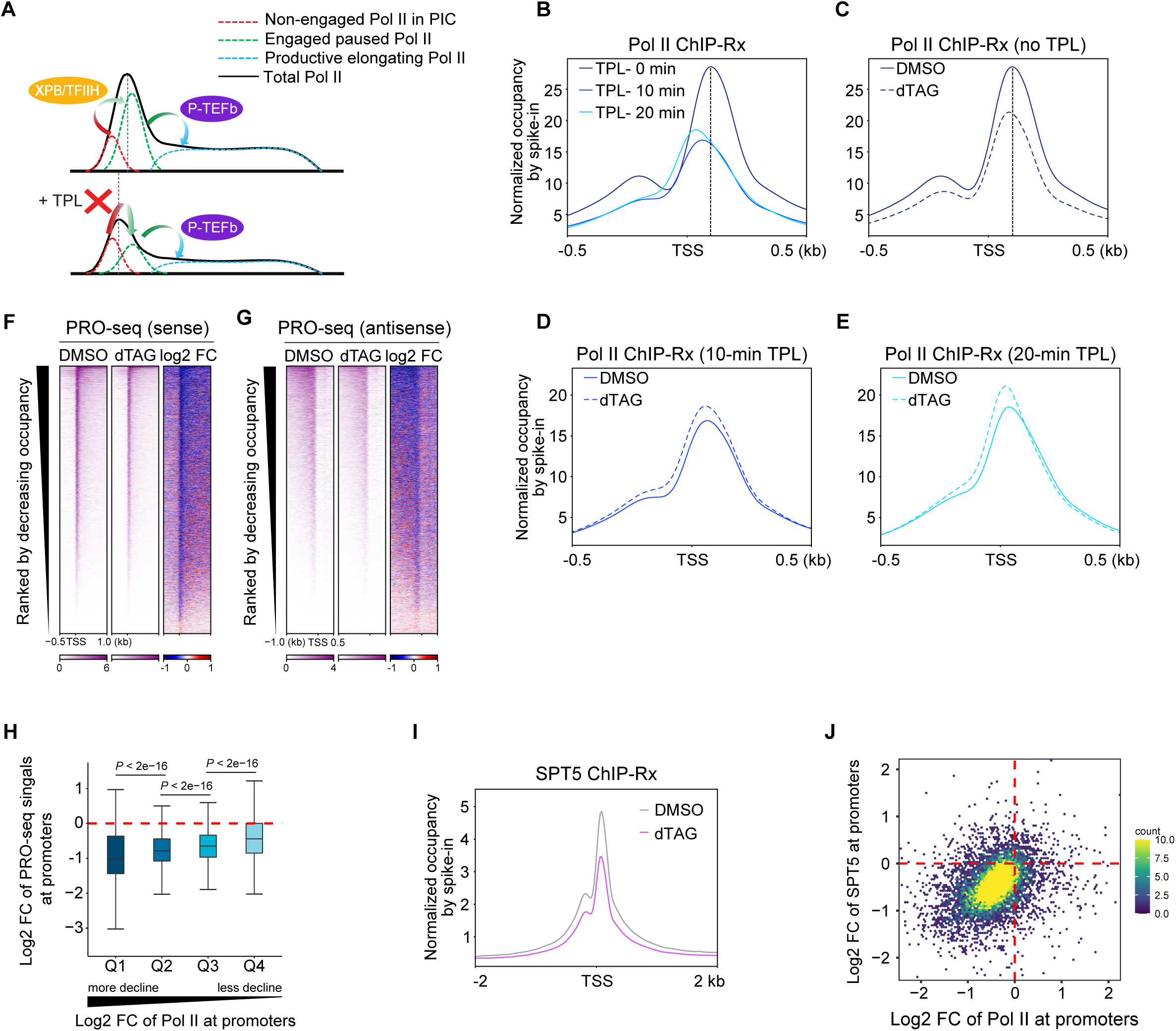
RBBP5 depletion and subsequent loss of H3K4me2 and H3K4me3 diminish the occupancy of paused but not initiating Pol II. (A) Schematic model for the composition of promoter-bound Pol II (top) and its occupancy shift upon TPL treatment (bottom). The promoter-bound Pol II (black line) is composed of nonengaged Pol II in PIC (red dash line) and engaged paused Pol II (green dash line). Upon TPL treatment, the transition from nonengaged Pol II to engaged Pol II is blocked, with pausing Pol II still releasing into gene bodies, causing the Pol II occupancy to shift upstream. (B) Metaplot showing the average level of Pol II occupancy in the DMSO condition of mRBBP5-dTAG mESCs upon TPL treatment. Metagenes are centered at the TSS of Pol II promoter peaks bound genes. (C-E) Metaplots showing the average level of Pol II occupancy centered at TSS of Pol II promoter peaks bound genes in mRBBP5-dTAG mESCs treated without TPL treatment (C) or treated TPL for 10 minutes (D) or 20 minutes (E). (F and G) Heatmaps of PRO-seq signals (RPM per bp and log2 fold change) for sense transcription (F) and antisense transcription (G) centered at the TSS of genes with H3K4me3 promoter peaks ranked by decreasing Pol II occupancy. (H) Boxplot showing the correlation of log2 fold change of PRO-seq signal (dTAG versus DMSO) and Pol II occupancy (dTAG versus DMSO) for four equal groups based on fold change of Pol II in mRBBP5-dTAG mESCs treated with DMSO or dTAG for 24 hours. (I) Metaplot showing the average level of SPT5 occupancy centered at the TSS of genes with H3K4me3 promoter peaks. (J) Two-dimensional density plot comparing log2 fold change of Pol II signal (x axis) and SPT5 signals (y axis) (dTAG versus DMSO) at H3K4me3 bound promoters in mRBBP5-dTAG cells. See also Figure S4.

To further distinguish Pol II in the PIC from Pol II in the paused state, we conducted spike-in normalized PRO-seq that specifically measures the engaged, paused polymerases but not the PIC form of Pol II. The level of PRO-seq signal decreases as profoundly as that of Pol II at promoters in RBBP5-depleted mESCs for both sense and antisense transcription (Fig. 4F-G, Fig. S4A-B). These changes in PRO-seq signal are correlated with overall Pol II levels at promoters, further supporting the notion that the decrease in Pol II at promoters results from loss of the pausing form of Pol II (Fig. 4H). Accordingly, the level of Pol II phosphorylated at serine 5 of CTD (pSer5), the form of polymerases in the pausing complex, is reduced in dTAG-treated RBBP5-dTAG mESCs (Fig. S4C).

Paused Pol II, but not initiating Pol II, is tightly associated with SPT5, with SPT5 both stabilizing the paused form of Pol II and subsequently traveling with Pol II after being released into productive elongation (*47, 73*). As measured by ChIP-Rx, the occupancy of SPT5 is diminished upon dTAG-induced RBBP5 degradation and reduction of H3K4me2, H3K4me3 and Pol II at promoters (Fig. 4I, Fig. S4D). As illustrated in the two-dimensional density plot, the reduced occupancy of SPT5 is overall proportional with that of Pol II at promoters, with most promoters showing reduced levels of both Pol II and SPT5 (Fig. 4J). These results collectively lead to the conclusion that loss of H3K4me2 and H3K4me3 compromises the maintenance of paused polymerases.

### Loss of H3K4 methylation compromises gene expression

We next sought to scrutinize the fate of the paused Pol II lost in response to RBBP5 depletion and the subsequent loss in H3K4 methylation. Plotting Pol II occupancy from TSS to TES reveals that RBBP5 degradation leads to decreased Pol II levels within gene bodies (Fig. 5A). A comparison of Pol II ChIP-Rx (Fig. 5B) and PRO-seq (Fig. 5C) data reveals that the decrease in Pol II occupancy at promoters is correlated with Pol II change in gene bodies. Consistently, genes with a greater decline of Pol II at promoters also have greater loss of Pol II in gene bodies (Fig. S5A-B). These results suggest that the observed loss of paused polymerases in dTAG-treated RBBP5-dTAG cells is not a result of Pol II being released into gene bodies for productive elongation, but instead that the paused Pol II was lost from chromatin.

**Figure 5.**
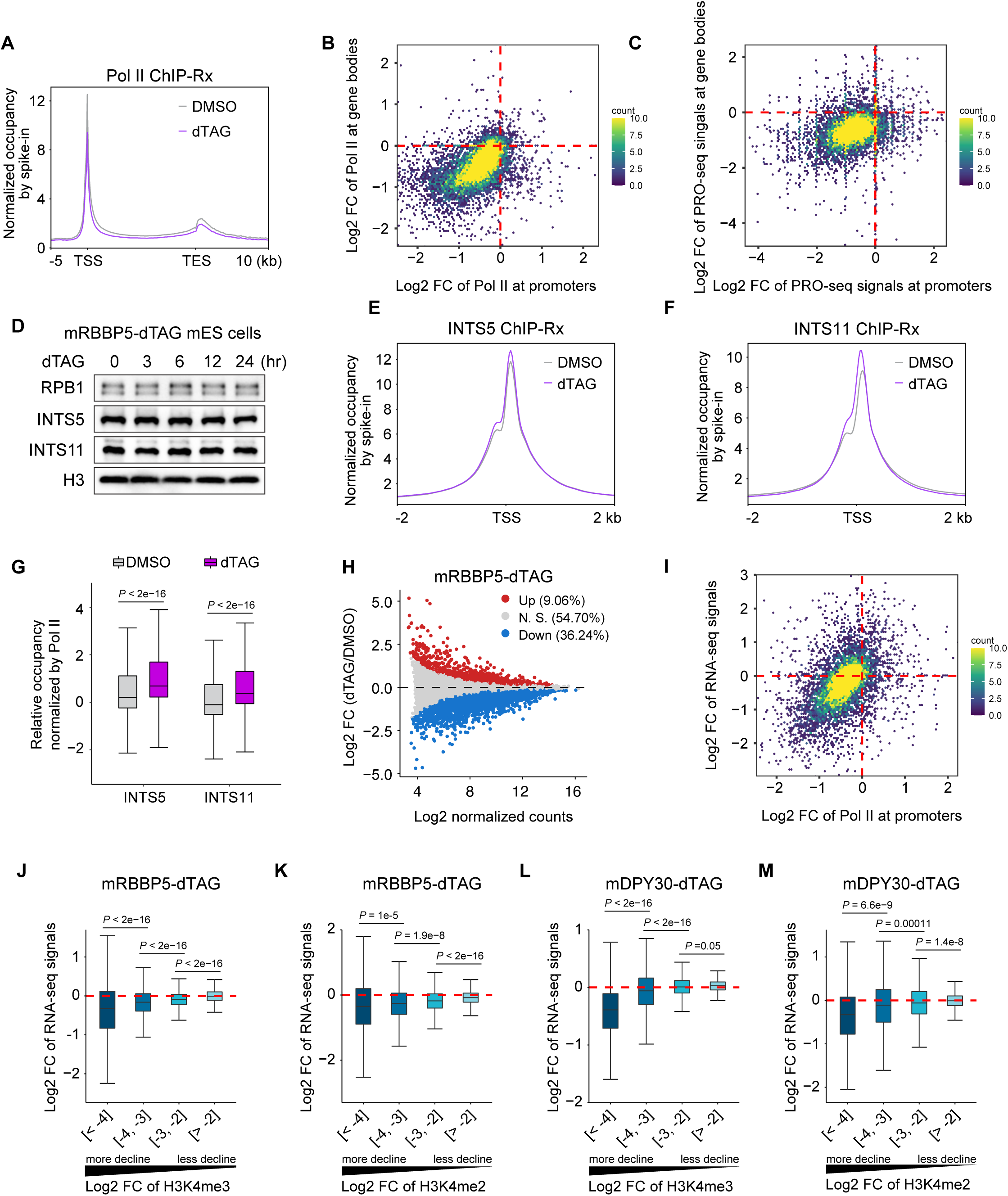
The loss of H3K4me2 and H3K4me3 compromise gene expression. (A) Metaplot showing the average Pol II occupancy genes with H3K4me3 promoter peaks as measured by ChIP-Rx in mRBBP5-dTAG cells treated with DMSO or dTAG. Pol II occupancy is represented by its largest subunit RPB1. (B and C) Two-dimensional density plot comparing the log2 fold change of Pol II ChIP-Rx signal (B) and PRO-seq signal (C) at promoters (x axis) and gene bodies (y axis) (dTAG versus DMSO) for genes with H3K4me3 promoter peaks. (D) Western blotting of mSPT5-dTAG mESCs with indicated time-course of dTAG treatment. (E and F) Metaplots showing the average level of INTS5 (E) and INTS11 (F) occupancies centered at the TSS of genes with H3K4me3 promoter peaks. (G) Boxplot showing the relative occupancies of INTS5 and INTS11 compared with Pol II in mRBBP5-dTAG mESCs treated with DMSO or dTAG. (H) M (log2 fold change, y axis) versus A (log2 average normalized read abundance, x axis) (MA) plot of spike-in normalized RNA-seq showing the change in gene expression by dTAG treatment for 24 hours in mRBBP5-dTAG cells. (I) Two-dimensional density plot comparing the log2 fold change of Pol II ChIP-Rx signal at promoters (x axis) and RNA-seq signal (y axis) (dTAG versus DMSO) for genes with H3K4me3 promoter peaks. (J and K) Boxplots showing the correlation of log2 fold change of RNA-seq singals (dTAG versus DMSO) and H3K4me3 (J) or H3K4me2 (K) (dTAG versus DMSO, grouped based on the declining extent of H3K4me3/H3K4me2 occupancy at promoters) in mRBBP5-dTAG mESCs. (L and M) Boxplots showing the correlation of log2 fold change of RNA-seq signal (dTAG versus DMSO) and H3K4me3 (L) or H3K4me2 (M) (dTAG versus DMSO, grouped based on the declining extent of H3K4me3/H3K4me2 occupancy at promoters) in mDPY30-dTAG mESCs. See also Figure S5.

The bulk levels of Pol II as assessed by western blotting remained unaltered following RBBP5 depletion, implying Pol II loss from chromatin is unlikely to be through polymerase protein degradation (Fig. 5D). Therefore, we surmised that the loss of paused Pol II upon RBBP5 and subsequent H3K4me2 and H3K4me3 loss could be due to premature termination of transcription, a pathway undergone by ∼80% of paused Pol II according to a recent study (*74*). As the dual-enzymatic Integrator-PP2A (INTAC) complex is mainly responsible for the premature termination at promoters, we measured its occupancy by performing ChIP-Rx of INTS5, a scaffold subunit of the “Shoulder” module that recruits the phosphatase module, and INTS11, the catalytic subunit of the RNA endonuclease module. In contrast to the reduction in levels of paused Pol II and its associating factor SPT5, the binding of INTS5 and INTS11 was observed to have a slight increase following RBBP5 depletion in mESCs (Fig. 5E-F, Fig. S5C-D). Because INTAC tightly associates with and depends on Pol II and SPT5 for its chromatin binding (*75–77*), we used promoter-bound Pol II and SPT5 to normalize the occupancy of INTAC. The relative occupancy of INTS5 and INTS11 exhibited a marked elevation upon RBBP5 depletion compared with Pol II (Fig. 5G) or SPT5 (Fig. S5E). These data support the conclusion that the fate of decreased paused polymerases upon loss of H3K4 methylation is mainly through premature termination, presumably catalyzed by INTAC.

We next conducted spike-in normalized RNA-seq to examine the impact of reduced levels of H3K4 methylation on gene expression. Upon RBBP5 degradation, we observed that the number of significantly downregulated genes was four times as many as significantly upregulated genes (Fig. 5H). Moreover, these changes in gene expression are highly correlated with the alterations in Pol II occupancy at promoters upon RBBP5 loss (Fig. 5I). Importantly, genes with the most severe decline in H3K4me2 and H3K4me3 levels also exhibit the greatest downregulation of gene expression (Fig. 5J-K).

To further validate the connection between H3K4 methylation and gene expression, we performed RNA-seq in DPY30-dTAG cells with 24-hour DMSO or dTAG treatment and analyzed its correlation with the loss of H3K4me2 and H3K4me3. Consistently, more genes are downregulated than upregulated (Fig. S5F). Furthermore, genes exhibiting a greater loss of H3K4 methylation had a more evident loss of gene expression (Fig. 5L-M). Together, these results reveal a dependency for H3K4 methylation in the optimal activation of transcription through regulation of paused Pol II.

## Discussion

Through the use of rapid degradation systems, our studies have revealed the dynamic deposition and removal of histone H3K4 methylation, as well as its unexpected role in modulating the stability of RNA polymerase II pausing. Rapid degradation of RBBP5 and DPY30, two shared subunits of all six COMPASS members in mammals, results in a remarkable decline in H3K4me2 and H3K4me3 by 6 hours and a near-complete loss by 12 to 24 hours. The removal of the higher H3K4 methylation states is dependent on the catalytic activity of KDM5 family of histone demethylases, as chemical inhibition of KDM5 blocks the fast turnover of H3K4 methylation upon COMPASS subunit degradation. Surprisingly, the near-complete loss of H3K4me2 and H3K4me3 had no major impact on the recruitment of the general transcription factor TFIID and initiating polymerases, a process that is generally thought to be promoted by the H3K4me3-TAF3 interaction. In contrast, the occupancy of the promoter-proximally pausing form of Pol II decreased pronouncedly following the removal of H3K4me2 and H3K4me3, which is likely due to premature termination of transcription by the INTAC complex, as the ratio of INTAC to Pol II occupancy is increased upon loss of H3K4me2 and H3K4me3. Consequently, levels of elongating polymerases and gene expression decline in cells with the ablation of H3K4me2 and H3K4me3.

Unlike histone acetylation, which has a half-life on the order of minutes (*78, 79*), histone methylation is believed to be more stable and can function as inheritable epigenetic marks (*17, 80, 81*). Although H3K4me2 and H3K4me3 are used as markers of active gene promoters, they are also found at transcriptionally poised genes (*82*), and are particularly pervasive over HOX gene clusters (*27*), where they are considered to antagonize epigenetic silencing (*82–86*). However, our understanding of the overall dynamics of H3K4me2 and H3K4me3 in mammals at a genome-wide scale is lacking, hindering us from exploring the regulatory roles of H3K4 methylation. Our finding that disruption of COMPASS promptly leads to a drop in H3K4me2 and H3K4me3 levels, reveals an unexpectedly high turnover rate of these modifications. Furthermore, it also suggests that H3K4me2 and H3K4me3 are constantly balanced by the competition between methyltransferases and demethylases. Therefore, COMPASS is continuously required to maintain global H3K4 methylation levels. This high turnover property would endow them with the capacity to modulate more dynamically regulated processes such as transcription.

Various roles for H3K4 methylation have been ascribed to a spectrum of reader proteins that function in various processes including transcription. Among them, TAF3, a key subunit of TFIID, arguably has the most intimate potential connection with active transcription. Our observation that an apparent near-complete loss of H3K4me2 and H3K4me3 had no noticeable impact on TFIID occupancy suggests that the role of H3K4me3 in transcription initiation needs further consideration. It is possible that the H3K4me3-TAF3 interaction could promote TFIID recruitment and PIC assembly at specific loci under certain biological contexts as previously demonstrated (*38, 39, 87*). For example, TFIID could be allosterically stimulated by TAF3 binding to H3K4me3 to promote the activation of transcription in response to specific signaling cues rather than reinitiation of already established transcribing genes, the latter of which would likely be the primary readout of our experimental perturbations.

Cryo-EM structures of the PIC in complex with the +1 nucleosome demonstrated extensive interactions between the PIC and the nucleosome core independently of histone tails (*88*). Moreover, TFIID contains other histone modification readers, such as TAF1 that recognizes histone acetylation through its bromodomains (*89, 90*). Together, these lines of evidence suggest a model whereby the nucleosome core and the panoply of modifications on the histone tails collectively regulate TFIID and PIC assembly on chromatin. Therefore, the loss of one histone modification could be buffered by other histone modifications, such that limited effects on transcription initiation could be observed under certain experimental conditions. Future studies simultaneously disrupting multiple histone modifications or by interfering with nucleosome assembly/disassembly are warranted to unveil the complex communication between the transcription machinery and nucleosome.

Paused RNA Pol II dwells at promoters for varying lengths of time, ranging from seconds to more than 10 minutes, implying that the stability of pausing is likely subject to dynamic regulation (*72, 74, 91-95*). Most of the mechanistic studies of pausing have been focused on several Pol II-bound pausing regulatory factors, including but not limited to NELF, DSIF, PAF1C and INTAC (*2, 46, 96-98*). Our findings lead us to propose a causal role of H3K4me2 and H3K4me3 in retaining paused Pol II at promoters. Importantly, the loss of the transcriptionally pausing form of Pol II following reductions in H3K4me2 and H3K4me3 levels is accompanied by a further decline in elongating polymerases and the subsequent downregulation of gene expression, suggesting that Pol II in the transcriptionally paused state is terminated prematurely rather than being released into gene bodies for productive transcription elongation. Supporting this notion, the loss of H3K4me2 and H3K4me3 induces a relative enrichment of promoter-bound INTAC complex, which has recently been found to be a major regulator of premature promoter-proximal termination in higher organisms (*75, 96, 98-106*). However, it is still unclear whether H3K4me2 and H3K4me3 regulate pausing directly through modulating chromatin structure or by recruiting effector proteins implicated in paused Pol II stability.

Interestingly, one of the core COMPASS subunits that are collectively known as WRAD, includes DPY30, which also happens to be a subunit of the nucleosome remodeling factor NURF, a chromatin remodeling complex that contains H3K4me3 readers BPTF and BAP18 (*107, 108*), raising the possibility of crosstalk between COMPASS and NURF in the regulation of transcription. Therefore, it will be particularly important to systematically evaluate the interactions of various H3K4 methylation readers with other transcription factors that impact Pol II pausing and its subsequent release into productive elongation. In conclusion, the studies presented here identify, while raising new questions about, a previously unrecognized interplay between histone modifying enzymes, chromatin structure, and the regulation of transcription in metazoans.

## Acknowledgments

This work was supported by grants from the National Key R&D Program of China (2021YFA1301700, 2021YFA1300100), the National Natural Science Foundation of China (32070636, 32270588), and the Shanghai Natural Science Foundation (20ZR1412100, 22ZR1412400).

## Author contributions

H.S., T.N., Q.Z. and W.Z. conducted the experiments. S.A. and P.L. performed the bioinformatics analyses. C.F.X and S.A. wrote the manuscript. C.F.X supervised the project.

## Declaration of interests

The authors declare no competing interests.

## Supplemental Figure Legends

**Figure S1. Related to Figure 1.**
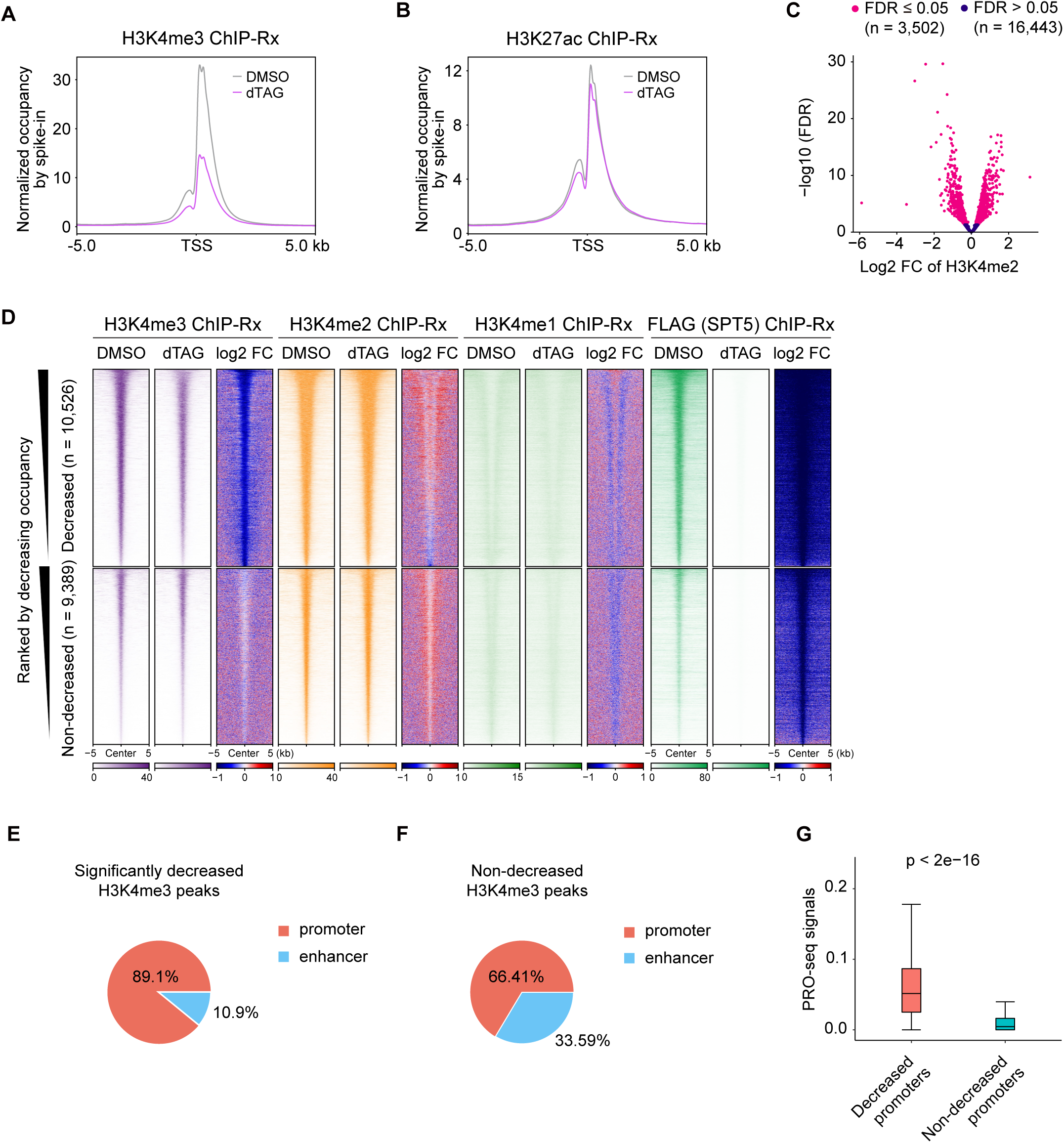
(A and B) Metaplots showing the average H3K4me3 (A) and H3K27ac (B) occupancies centered at TSS measured by ChIP-Rx in hSPT5-dTAG DLD-1 cells treated with DMSO or dTAG. (C) Volcano plot showing the log2 fold change of H3K4me2 peaks of 6-hour dTAG versus DMSO treatment. (D) Heatmaps showing the occupancies of decreased and non-decreased groups of H3K4me3, H3K4me2, H3K4me1, and FLAG (SPT5) (RPM per bp and log2 fold change) in mSPT5-dTAG mESCs centered at H3K4me3 peak centers ranked by decreasing H3K4me3 occupancy in the DMSO condition. (E and F) Genomic distribution of the significantly decreased (E) and non-decreased (F) H3K4me3 peaks. (G) Quantification of the average PRO-seq signal of H3K4me3 peaks bound PRO-seq promoters in the DMSO condition in mSPT5-dTAG cells.

**Figure S2. Related to Figure 2.**
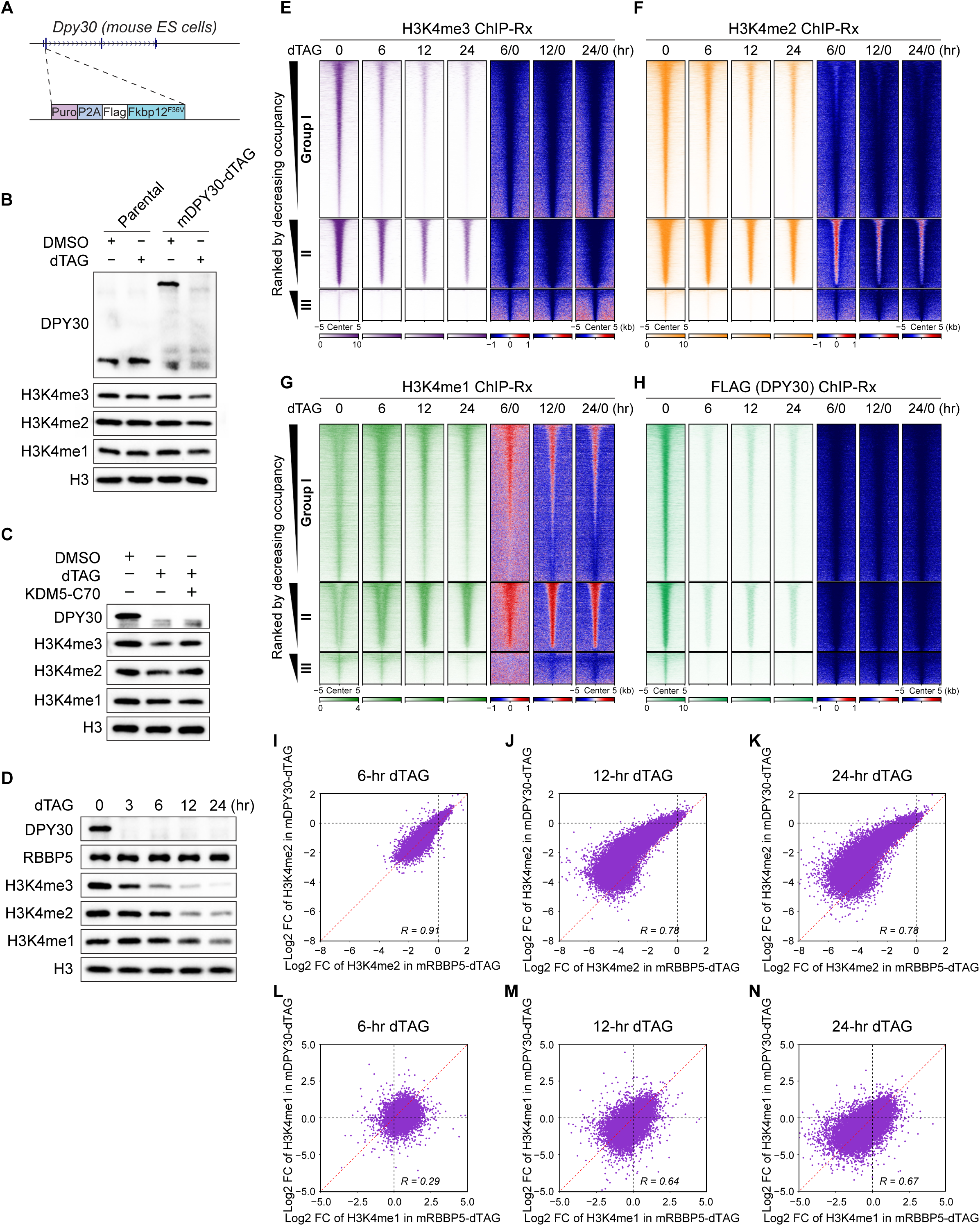
(A) Schematic of the generation of mDPY30-dTAG mESCs by knock-in. Western blotting of whole-cell extracts of parental and mDPY30-dTAG mESCs treated with DMSO or dTAG for 6 hours. H3 is a loading control. (C) Western blotting of dTAG treatment with or without pan-KDM5 inhibition by KDM5-C70 in mDPY30-dTAG mESCs. (D) Western blotting of mDPY30-dTAG mESCs with time-course dTAG treatment. (E-H) Heatmaps showing three groups of H3K4me3, H3K4me2, and H3K4me1 occupancy (RPM per bp and log2 fold change) ranked by decreasing H3K4me3 occupancy in the DMSO condition in mDPY30-dTAG mESCs. The peaks are centered at the H3K4me3 peak centers. (I-K) Scatterplots depicting the correlations of log2 fold change of H3K4me2 peaks of 6- (I), 12- (J), and 24-hour dTAG (K) versus DMSO treatment between mRBBP5-dTAG cells and mDPY30-dTAG cells. (L-N) Scatterplots depicting the correlations of log2 fold change of H3K4me1 peaks of 6- (L), 12- (M), and 24-hour dTAG (N) versus DMSO treatment between mRBBP5-dTAG cells and mDPY30-dTAG cells.

**Figure S3. Related to Figure 3.**
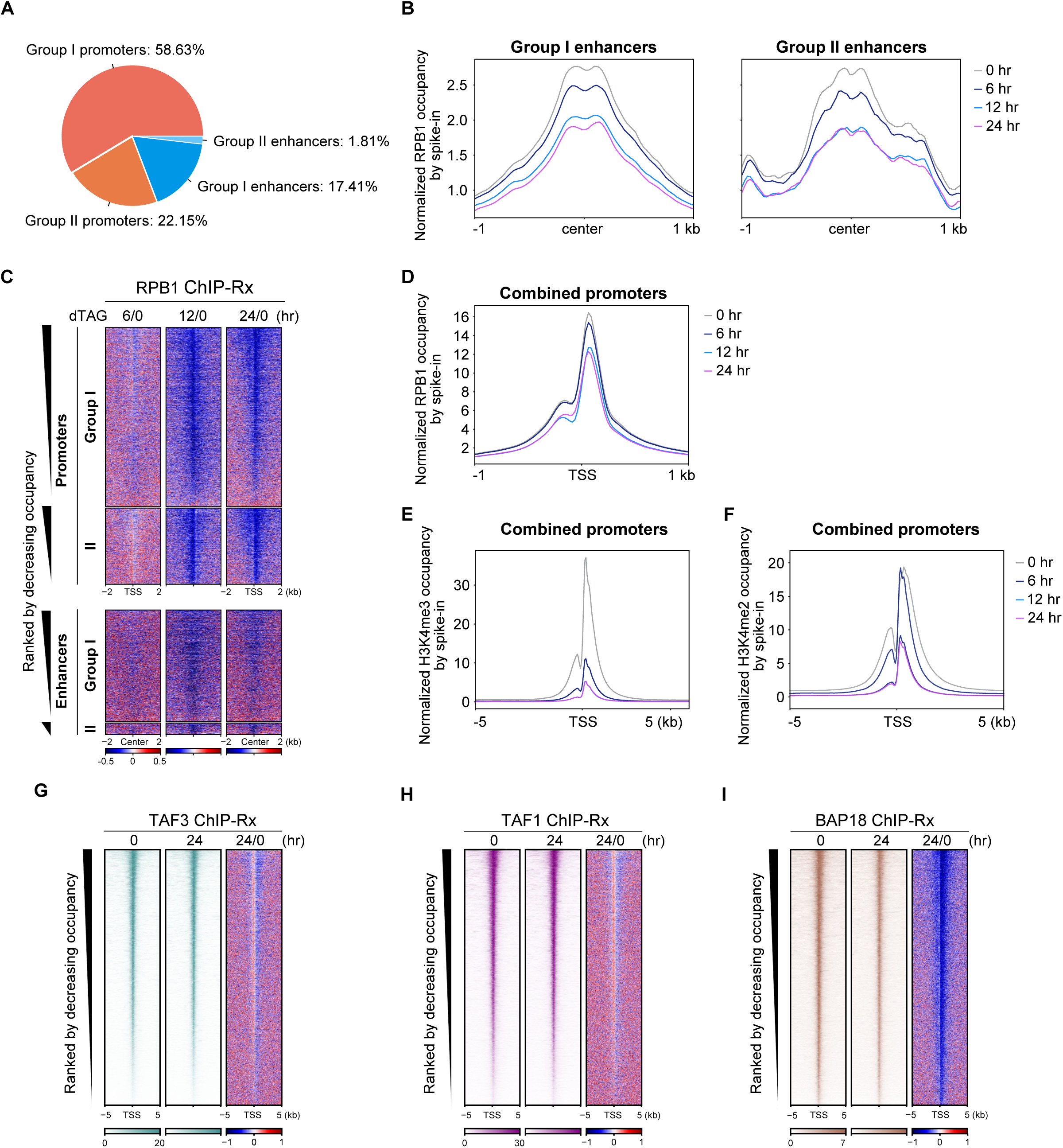
(A) Genomic distribution of Group I and Group II H3K4me3 peaks. (B) Metaplots showing the average RPB1 occupancy at Group I and Group II enhancers centered at H3K4me3 peak centers measured by ChIP-Rx in mRBBP5-dTAG mESCs treated with dTAG for 0, 6, 12, and 24 hours. (C) Heatmaps showing log2 fold change of Pol II signal (dTAG versus DMSO) in mRBBP5-dTAG cells. (D-F) Metaplots showing the average RPB1 (D), H3K4me3 (E), and H3K4me2 (F) occupancies at combined promoters centered at the TSS of H3K4me3 promoter peaks bound genes measured by ChIP-Rx in mRBBP5-dTAG mESCs treated with dTAG for 0, 6, 12, and 24 hours. (G-I) Heatmaps of TAF3 (G), TAF1 (H), and BAP18 (I) occupancies (RPM per bp and log2 fold change) centered at the TSS of genes with H3K4me3 promoter peaks.

**Figure S4. Related to Figure 4.**
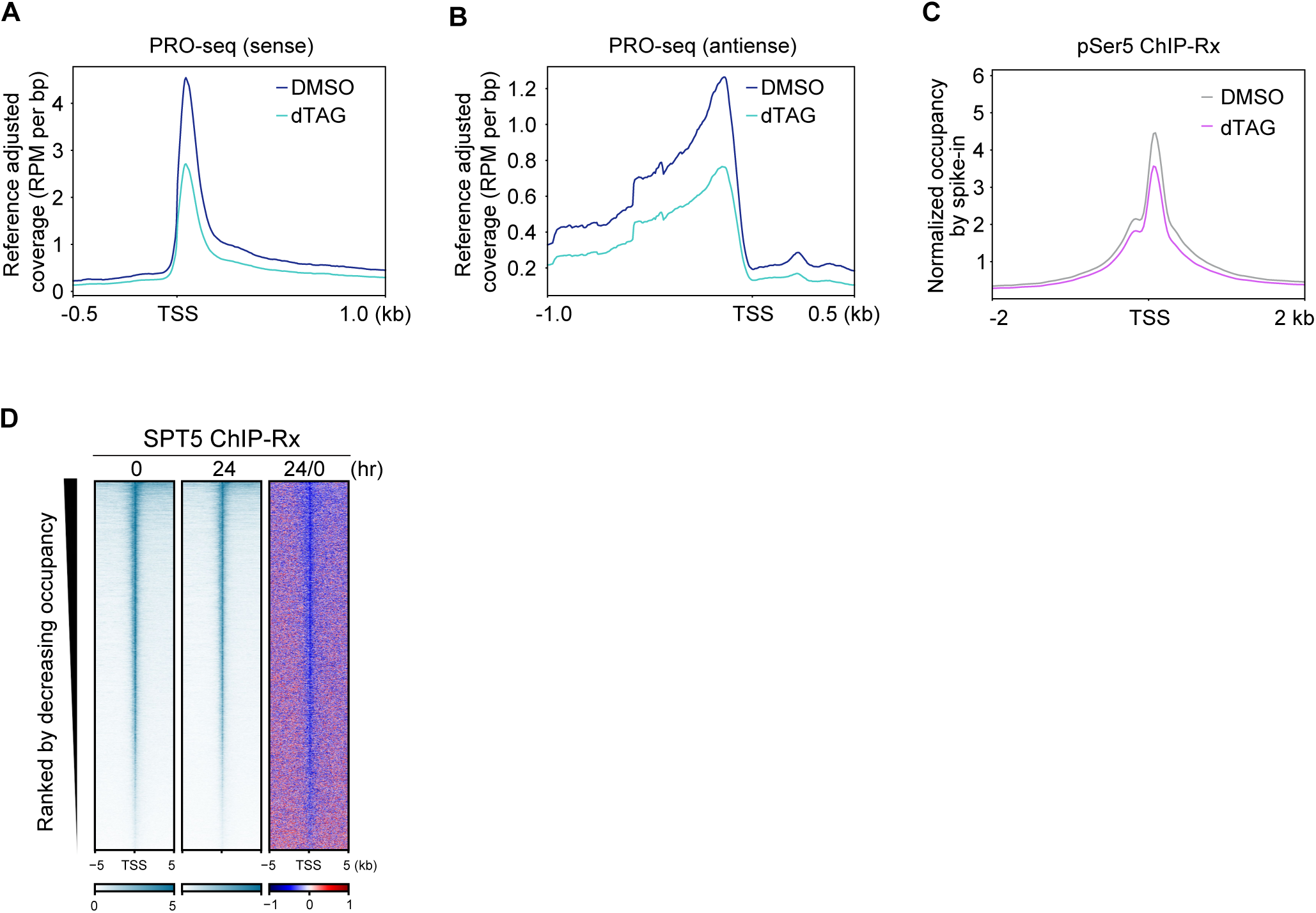
(A and B) Metaplot profiles of PRO-seq signal showing the average signals of sense (A) and antisense (B) transcription in DMSO- or dTAG-treated mRBBP5-dTAG mESCs. (C) Metaplot showing the average occupancy of Pol II pSer5. (D) Heatmaps of SPT5 occupancy (RPM per bp and log2 fold change) centered at the TSS of genes with H3K4me3 promoter peaks.

**Figure S5. Related to Figure 5.**
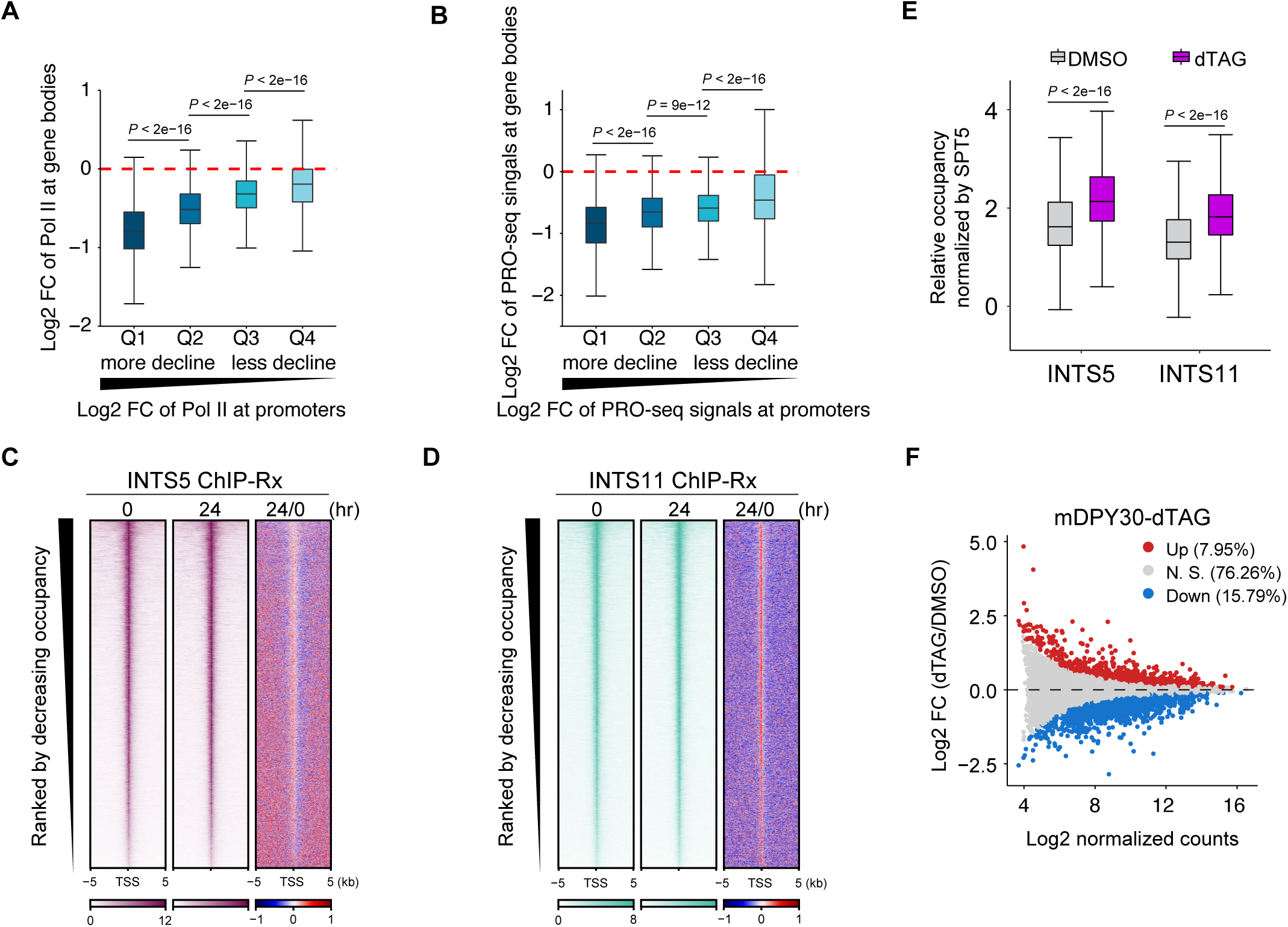
(A) Boxplots showing the correlation of the log2 fold change of Pol II at promoters and gene bodies (dTAG versus DMSO) for four equal groups based on the fold change of Pol II at promoters in mRBBP5-dTAG mESCs. (B) Boxplots showing the correlation of the log2 fold change of PRO-seq signal at promoters and gene bodies (dTAG versus DMSO) for four equal groups based on the fold change of PRO-seq signal at promoters in mRBBP5-dTAG mESCs. (C and D) Heatmaps of INTS5 (C) and INTS11 (D) occupancies (RPM per bp and log2 fold change) centered at the TSS of genes with H3K4me3 promoter peaks. (E) Boxplots showing the relative occupancies of INTS5 and INTS11 compared with SPT5 in mRBBP5-dTAG mESCs treated with DMSO or dTAG. (F) MA plot of spike-in normalized RNA-seq showing the gene expression changes by dTAG treatment for 24 hours in mDPY30-dTAG cells.

## METHODS

### Genome editing for endogenous knock-in dTAG cells

Endogenous knock-in of dTAG was conducted following previously described methods (*109*). To generate endogenous knock-in DLD-1 cells, PITCh cutting sgRNA and donor plasmid were mixed with 1 × 10^6^ DLD-1 cells followed by electroporation (Neon). Cells were immediately transferred to 6-well plates containing DMEM media without antibiotics. After recovery for 2 days, cells were diluted and cultured with 1μg/ml puromycin (Meilunbio) for 10-14 days. The surviving clones were picked, transferred to 96-well plates and expanded before genotyping by PCR. Protein degradation efficiencies of successful homogeneous clones were verified by dTAG-13 treatment for 3 hours followed by western blotting.

To generate endogenous knock-in mouse ES cells (mESCs), PITCh plasmids, sgRNA plasmids and donor plasmids were transfected with Lipo3000 Reagents (Invitrogen) according to the manufacturer’s instruction. 5 × 10^5^ cells were seeded into 6-well plates one day in advance. 1 μg sgRNA-Cas9 plasmids, 1μg donor plasmids and 0.5 μg PITCh plasmids were mixed with P3000 and Lipo3000 in Opti-MEM. The mixture was incubated for 15 min at room temperature, and then added to the cell culture and incubated with cells overnight at 37 ℃. Cells were trypsinized and seeded in 10 cm dishes with 1 μg/ml puromycin (Meilunbio) for 10-14 days. Single-clone colony picking and clone validation were performed as described above.

### Chromatin IP with reference exogenous genome (ChIP-Rx)

ChIP-Rx was performed as described previously (*49*). 5 × 10^7^ mESCs were crosslinked with 1% formaldehyde (Sigma) for 10 min at room temperature. The reaction was quenched by adding glycine to a final concentration of 125 mM and incubating for 5min. Cells were washed in cold PBS, resuspended in lysis buffer (50 mM HEPES pH7.4, 150 mM NaCl, 2 mM EDTA, 0.1% SDS, 0.1% sodium deoxycholate, 1 × protease inhibitor (Roche)). Fixed cells were sonicated (Qsonica) to obtain chromatin fragments between 200-700bp, and centrifuged to collect the soluble fraction. Human DLD-1 cells were used as a spike-in for normalization. Mixed lysates were then incubated overnight with antibodies and magnetic Protein A/G beads for 2 hours. The beads were washed 3 times with High Salt Wash buffer (20 mM HEPES pH7.4, 500 mM NaCl, 1 mM EDTA, 1.0% NP-40, 0.25% sodium deoxycholate), twice wiht Low Salt Wash buffer (20 mM HEPES pH 7.4, 150 mM NaCl, 1 mM EDTA, 0.5% NP-40, 0.1% sodium deoxycholate), and once with TE buffer containing 50 mM NaCl. Beads were eluded with Elution buffer (50 mM Tris-HCl pH 8.0, 10 mM EDTA, 1.0% SDS). The eluted samples were digested with Proteinase K and purified by Phenol/Chloroform/Isoamyl Alcohol extraction. The sequencing libraries were prepared with VAHTS Universal Plus DNA Library Prep Kit for Illumina (Vazymes). Libraries were sequenced using an Illumina HiSeq X Ten or NovaSeq 6000 platform (Annoroad Gene Technology, Beijing, China).

### Transient transcriptome sequencing (TT-seq)

TT-seq was carried out as described previously (*110*). mESCs were grown in 15 cm dishes, and the RNA was labelled with 4-thiouridine (4sU) (Cool Chemistry) in vivo for 15 min. TRIzol (Invitrogen) was added to stop the reaction and total RNA was extracted according to instructions. As a control, 4sU-labeled DLD-1 RNA was mixed as the spike-in. The mix was fragmented with 0.2 M NaOH for 14min and Tris-HCl (pH 6.8) was added to stop the fragmentation. Biotinylation of labeled RNA was carried out in 150 μL of biotinylation mix (100 μg fragmented total RNA, 10 mM HEPES pH 7.5, 1 mM EDTA, 0.167 mg/mL MTSEA-biotin) for 30 min and purified with Streptavidin Magnetic Beads (NEB). The biotinylated RNA was eluted with 100 mM DTT and purified with RNA Clean beads (Vazymes). The RNA libraries were constructed from the 200 μg purified RNA sample using Stranded mRNA-seq Lib Prep Module for Illumina (ABclonal, RK20349) following the guidelines of the manufacturers and sequenced with a NovaSeq 6000 platform (Annoroad Gene Technology, Beijing, China).

### Precision run-on sequencing (PRO-seq)

PRO-seq was performed according to the previously published protocol with minor modifications (*110, 111*). mESCs were rinsed twice with 5 ml of cold 1 × PBS and scraped with 5ml permeabilization buffer (10 mM Tris-HCl pH 8.0, 5% glycerol, 250 mM sucrose, 10 mM KCl, 5 mM MgCl_2_, 1 mM EGTA, 0.5 mM DTT, 0.1% Igepal, 0.05% Tween-20, protease inhibitors cocktail (Roche), 4 U/mL RNase inhibitor (SUPERaseIN)). The resuspended cells were incubated for up to 5min on ice, then transferred for centrifugation and washing in ice-cold wash buffer (10 mM Tris-HCl pH 8.0, 10 mM KCl, 5% glycerol, 5 mM MgCl2, 0.5 mM DTT, 4 U/mL RNase inhibitor). Permeabilized cells were resuspended in freezing buffer (50 mM Tris-HCl pH 8.0, 40% glycerol, 5 mM MgCl2, 1 mM EDTA, 0.5 mM DTT, 4 U/mL RNase inhibitor) and immediately frozen in liquid nitrogen. The permeabilized cells were stored at −80 ℃ until usage.

Permeabilized cells were mixed with spike-in DLD-1 cells. Nuclear run-on reactions were performed with 2 × nuclear run-on reaction mixture (10 mM Tris-HCl pH 8.0, 300 mM KCl, 1% Sarkosyl, 5 mM MgCl2, 1 mM DTT, 40 μM Biotin-11-A/C/G/UTP (Perkin-Elmer), 0.8 U/mL RNase inhibitor) and incubated for 5min at 37℃. Nascent RNA was extracted by Trizol LS Reagent (Invitrogen). RNA fragmentation was performed with 0.25N NaOH on ice for 10 min and neutralized by adding 1M Tris-HCl pH6.8, followed by passing through a calibrated RNase-free P-30 column (Bio-Rad). After 3’ RNA adaptor ligation, RNA was purified by Streptavidin Magnetic Beads (NEB) in Binding buffer (10 mM Tris-HCl pH 7.4, 300 mM NaCl, 0.1% Triton X-100, 1 mM EDTA). The beads were washed once with High Salt Wash buffer (50 mM Tris-HCl pH 7.4, 2 M NaCl, 0.5% Triton X-100,1 mM EDTA) and once with Low Salt Wash buffer (5 mM Tris-HCl pH 7.4, 0.1% Triton X-100, 1 mM EDTA). On-Bead RNA 5’ hydroxyl repair was then performed with PNK mix (1 × PNK Buffer, 1 mM ATP, 10 U PNK) at 37 ℃ for 30 min. The RNA 5’ decapping was performed with RppH mix (1 × ThermoPol Buffer, 5 U RppH) at 37 ℃ for 1 h. On-Bead 5’ RNA adaptor ligation was performed by the ligation mix (1 × T4 RNA ligase buffer, 1 mM ATP, 15% PEG 8000, 10 U T4 RNA Ligase I) at 25 ℃ for 1 hour, followed by TRIzol elution of the adaptor ligated RNA. The RNA was reverse transcribed by the Maxima H Minus RT enzyme (Invitrogen). Full-scale libraries were amplified with Q5 enzyme mix (NEB). Libraries were sequenced with a NovaSeq 6000 platform (Annoroad Gene Technology, Beijing, China).

### Quantification and statistical analysis

#### Identification of transcription start sites (TSS) and enhancers

The identification of TSS in DLD-1 cells was performed using published PRO-cap bigwig files as described previously (*47, 112*). In brief, we collected the transcripts of known RefSeq protein-coding genes (*113*) and selected the transcript with the maximum PRO-cap signal at TSS – 10 bp and TSS + 300 bp regions of each gene as the representative transcript. Then the position of the maximum PRO-cap signal was defined as the TSS of the corresponding gene.

For mouse embryonic stem cells, we chose the longest transcript isoform for alternatively spliced protein-coding genes as the representative transcript. The chosen transcript and its TSS were then used for further analyses. For ChIP-Rx data, we defined the promoter region as TSS – 1000 bp to TSS + 1000 bp, the genebody region as TSS + 3000 bp to TES, and the intergenic enhancer region as from 2kb upstream and downstream of the protein-coding genes. For PRO-seq data, we defined the promoter region as TSS – 10 bp to TSS + 300 bp and the genebody region as TSS + 300 bp to TES.

#### ChIP-Rx data analysis

The raw ChIP-Rx reads were trimmed by Trim Galore v0.6.6 (https://www.bioinformatics.babraham.ac.uk/projects/trim_galore/) and aligned to the human hg19 and mouse mm10 assemblies using Bowtie v2.3.5.1 with default parameters (*114*). Low mapping quality reads (MAPQ < 30) and PCR duplicates were removed using SAMtools v1.9 (*115*) and Picard v2.23.3 (https://broadinstitute.github.io/picard/). We then collected the spike-in read number for each of the ChIP-Rx samples with SAMtools v1.9 (*115*) and generated the normalization factor as 1e6/spike-in_count. Normalized bigwig files were generated by deeptools v3.5.0 (*116*). The ENCODE blacklist regions were removed using bedtools v2.29.2 (*117*). Peak calling was performed by macs2 v2.2.6 with a q-value threshold of 0.05 (*118*). DiffBind R package v2.16.2 (*119*) was used to identify differential binding peaks.

#### PRO-seq data analysis

The paired PRO-seq reads were trimmed by Trim Galore v0.6.6 (https://www.bioinformatics.babraham.ac.uk/projects/trim_galore/) with read length >15 bp. After removal of molecular barcode (UMI) with fastp v0.21.0 (*120*), ribosomal RNA reads were discarded using Bowtie v2.3.5.1 with ‘‘--un-conc-gz’’ (*114*). Then the remaining reads were aligned to the hg19 or mm10 genome using Bowtie v2.3.5.1 with ‘‘--local --sensitive-local’’ (*114*). The mapped data were next deduplicated with UMI-tools to remove PCR duplicates based on the UMI sequences (*121*). We generated the normalization factor as described in ChIP-Rx and built strand-specific coverage tracks with deeptools v3.5.0 (*116*).

#### RNA-seq data analysis

Raw reads were trimmed as described for ChIP-Rx, followed by mapping to the hg19 or mm10 genome using STAR v2.7.5c with parameter ‘‘--outSAMtype BAM SortedByCoordinate --twopassMode Basic -- outFilterMismatchNmax 3’’ (*122*). Low mapping quality reads and PCR duplicates were removed using SAMtools v1.9 (*115*) and Picard v2.23.3 (https://broadinstitute.github.io/picard/). Calculation of the normalization factor was performed in the same way as described in ChIP-Rx. Strand-specific normalized bigwigs were generated by deeptools v3.5.0 (*116*). To identify differentially expressed genes, we collected the counts per gene by the featureCounts tool from Rsubread R package v2.0.1 (*123*). Differential expression analysis was then performed using DESeq2 R package v1.26.0 (*124*).

## Notes

### Competing Interest Statement

The authors have declared no competing interest.

